# Estimating the mean: behavioral and neural correlates of summary representations for time intervals

**DOI:** 10.1101/2024.10.08.617175

**Authors:** Taku Otsuka, Hakan Karsilar, Hedderik van Rijn

**Affiliations:** Experimental Psychology, University of Groningen, Groningen, 9712 TS, The Netherlands; Department of Life Sciences, The University of Tokyo, Tokyo, Meguro-ku, Komaba, 3-8-1, 153-8902, Japan

**Keywords:** time perception, summary representation, mean, distribution, CNV, P2

## Abstract

Our behavior is guided by the statistical regularities in the environment. Prior research on temporal context effects has demonstrated the dynamic processes through which humans adapt to the environment’s temporal regularities. However, learning temporal regularities not only entails dynamic adaptation to traces of previous individual events but also often requires the extraction and retention of summary statistics (e.g., the mean) of temporal distributions. To investigate these summary representations for temporal distributions and to test their sensitivity to distributional changes, we explicitly asked participants to extract the mean of different distributions of time intervals, which shared the same mean but varied in their variability specifically operationalized by the width and presentation frequency of the intervals. Our findings showed that the variability of the estimated mean increased with the distributions’ variability, even though the actual mean remained constant. We further examined how such learning of temporal distributions modulates EEG signals during subsequent temporal judgments. Analysis revealed that the contingent negative variation (CNV), predictive of single-trial RTs, was correlated with how much individuals’ estimates of the mean were affected by the distributions’ variability. Conversely, the post-interval P2 was not modulated by the distributions but predicted participants’ responses, suggesting that P2 reflects the perceived duration of an interval. Taken together, our results demonstrate not only that humans can accurately estimate the mean of a temporal distribution, but also that the representation of the mean becomes more uncertain as the variability of the distribution increases, as reflected neurally in the preparation-related CNV during temporal decisions.

## INTRODUCTION

From perception to decision-making to complex motor skills, our behavior is guided by the temporal structure of our surroundings. Understanding how our brain represents the temporal regularities of events is therefore a primary goal of psychology and neuroscience. Substantial evidence has shown that temporal context, both at the global (e.g., summary statistics of stimuli) and the local scales (e.g., presentation sequence of stimuli), affects temporal estimation (Cicchini et al., 2012; de Jong et al., 2021; Glasauer & Shi, 2021; Jazayeri & Shadlen, 2010; Jones & Mcauley, 2005; Lejeune & Wearden, 2009; Rhodes et al., 2023; Shi et al., 2013; Taatgen & van Rijn, 2011; Wang et al., 2023; Wiener et al., 2014). For instance, the central tendency effect in interval timing suggests that estimates of the current stimulus duration regress toward the mean of previously encountered stimuli (Vierordt, 1868). Importantly, these temporal context effects highlight an implicit and dynamic process of adapting to temporal regularities. More specifically, regardless of whether the effect is primarily driven by the global or local context, traces of previous stimuli modulate the temporal estimation of the immediately following stimulus on a trial-by-trial basis (de Jong et al., 2021; Taatgen & van Rijn, 2011; Togoli et al., 2021), and this effect persists regardless of explicit cues or instructions (Maaß et al., 2019; Roach et al., 2017)

However, in real-world scenarios, learning temporal regularities involves not only the implicit and dynamic adaptation to a series of previous events but also often requires the extraction and retention of global features of temporal distributions to guide future behavior. For example, in baseball, the throwing speed of the ball typically follows a unimodal distribution, similar to a Gaussian (MLB.com, 2024). In such cases, it is beneficial for a batter to extract the mean speed of previously pitched balls and the variability around this mean. This way, in the next game, the batter can predict the throwing speed of the same pitcher from the very first pitch, even without having observed any sample in the current game. Estimating the mean allows the batter to predict the most likely throwing speed (in the case of a unimodal distribution) while knowing the variability helps assess the reliability of the prediction. This example illustrates that, adaptive behavior likely requires more than just passive adaptation to statistical regularities; it also necessitates the active extraction of summary statistics.

The fundamental question then remains: How do humans extract and retain the summary statistics of temporal distributions? On the one hand, prior work on humans’ internal time scale explicitly instructed participants to estimate the mean of sample durations (Curtis & Rule, 1977; Ren et al., 2020; Schweickert et al., 2014; Wearden & Jones, 2007). While these studies indeed showed that participants were able to extract the mean of a range of time intervals, it is unclear whether the participants also encoded other features of these samples, such as their variability. On the other hand, studies on the aforementioned temporal context effects can also provide some clues. Context effects are known to align with the framework of Bayesian integration (Jazayeri & Shadlen, 2010; Petzschner et al., 2015; Sadibolova & Terhune, 2022; Shi et al., 2013; Shi & Burr, 2016), where the brain integrates the current sensory input (likelihood function) with the internal representation of the statistical regularities (prior distribution) to achieve optimal estimates (Knill & Pouget, 2004; Körding & Wolpert, 2006). Notably, using a model-based reconstruction of the prior distribution, Acerbi et al. (2012) suggested that humans can represent up to the third-order moments of a temporal distribution as an internal prior (i.e., mean, variance, and skewness). However, this reconstruction was based on temporal reproduction tasks, in which participants were asked to reproduce intervals coming from series of stimuli. Although such tasks can show that temporal estimates are influenced by the summary statistics of the underlying distribution, the effects are intrinsically passive and thus not sufficient to conclude that humans extract and retain these summary statistics. It can be argued that, once the dynamic adaptation to the temporal context has ended (i.e., after completing the reproduction task), the representations of the temporal regularities are entirely lost. Therefore, how sensitive humans are to subtle changes in learned temporal distributions when they need to extract summary statistics of the distribution remains an open question.

To this end, we explicitly asked participants to extract the mean of temporal distributions and examined whether the distributions’ variability affected their estimates of the mean. We developed a task consisting of two phases. In the first phase, participants performed an interval reproduction task in which they reproduced time intervals sampled from one of two Gaussian-like distributions with the same mean but different variability, operationalized by the width and/or frequency. In the second phase, they performed a temporal generalization task where they judged whether the test intervals matched the mean of the distribution encountered in the first phase.

Using the same task procedure, we further investigated whether the learning of temporal distributions modulates neural activities during later temporal judgments. EEG signatures in timing tasks such as the contingent negative variation (CNV) and the offset P2 have been linked to temporal context effects in humans (Baykan et al., 2023; Damsma et al., 2021; Wiener & Thompson, 2015). Research increasingly suggests that CNV is related to the preparation and anticipation of an upcoming event or action (Boehm et al., 2014; Breska & Deouell, 2014, 2017; Leuthold et al., 2004; Mento, 2013; Praamstra et al., 2006; Scheibe et al., 2010), while offset P2 is linked to the perceptual aspect of the current duration (Damsma et al., 2021; Kononowicz & van Rijn, 2014; Kruijne et al., 2021). We hypothesized that prior learning of temporal distributions would modulate the preparation state during later temporal decisions, but not the perceived time of stimuli. Therefore, we predicted that the CNV, but not the offset P2, would be modulated by the learning.

By capitalizing on the subjective estimates of the distribution’s mean and its associated uncertainty, we reveal that the summary representation of the mean of a temporal distribution is influenced by the distribution’s variability. Furthermore, we show that the effect of prior learning of temporal distributions is reflected in the preparation-related CNV during subsequent temporal judgments, but not in the offset P2, which instead indexes the perceived time of a stimulus.

## METHODS

### Experiment 1 (Behavioral)

#### Participants

Ninety first-year psychology students at the University of Groningen participated in the experiment for partial course credit. 30 participants were included in Experiment 1A (mean age = 20.7 years; 25 female), 30 in Experiment 1B (mean age = 19.6 years; 28 female), and 30 in Experiment 1C (mean age = 20.3 years; 24 female). We determined our sample size to be approximately double that of previous similar behavioral experiments to increase statistical power (Ren et al., 2020: 16 participants; Schweickert et al., 2014: 15 participants; Wearden & Jones, 2007: 16 participants). All participants reported normal or corrected-to-normal vision, normal hearing, and no history of psychiatric or neurological conditions. The experiment was approved by the Ethical Committee of the Faculty of Behavioural and Social Sciences (PSY-2324-S-0116). Written informed consent was obtained before the experiment.

#### Apparatus and stimuli

The experiment was programmed using OpenSesame 4.0 (Mathôt et al., 2012) with the Expyriment backend (Krause & Lindemann, 2014). Auditory stimuli were 440 Hz sine wave tones presented on Sennheiser HD 400S stereo headphones at a comfortable sound level. Visual stimuli were presented using a 27-inch Iiyama ProLite G27773HS monitor with a 1920 × 1080 resolution at 100 Hz. The left and right index-finger trigger buttons of a gamepad (SideWinder Plug & Play Game Pad, Microsoft) were used to record responses. Participants sat in a dimly lit room approximately 60 cm from the screen. A gray background was maintained during the entire experiment. A fixation dot (1.7°in diameter) was presented in the center of the screen. A dark gray square, circle, or star was presented in a 3.5 × 3.5°area in the top left of the screen during the entire task (see Fig. 1A).

**Figure 1.**
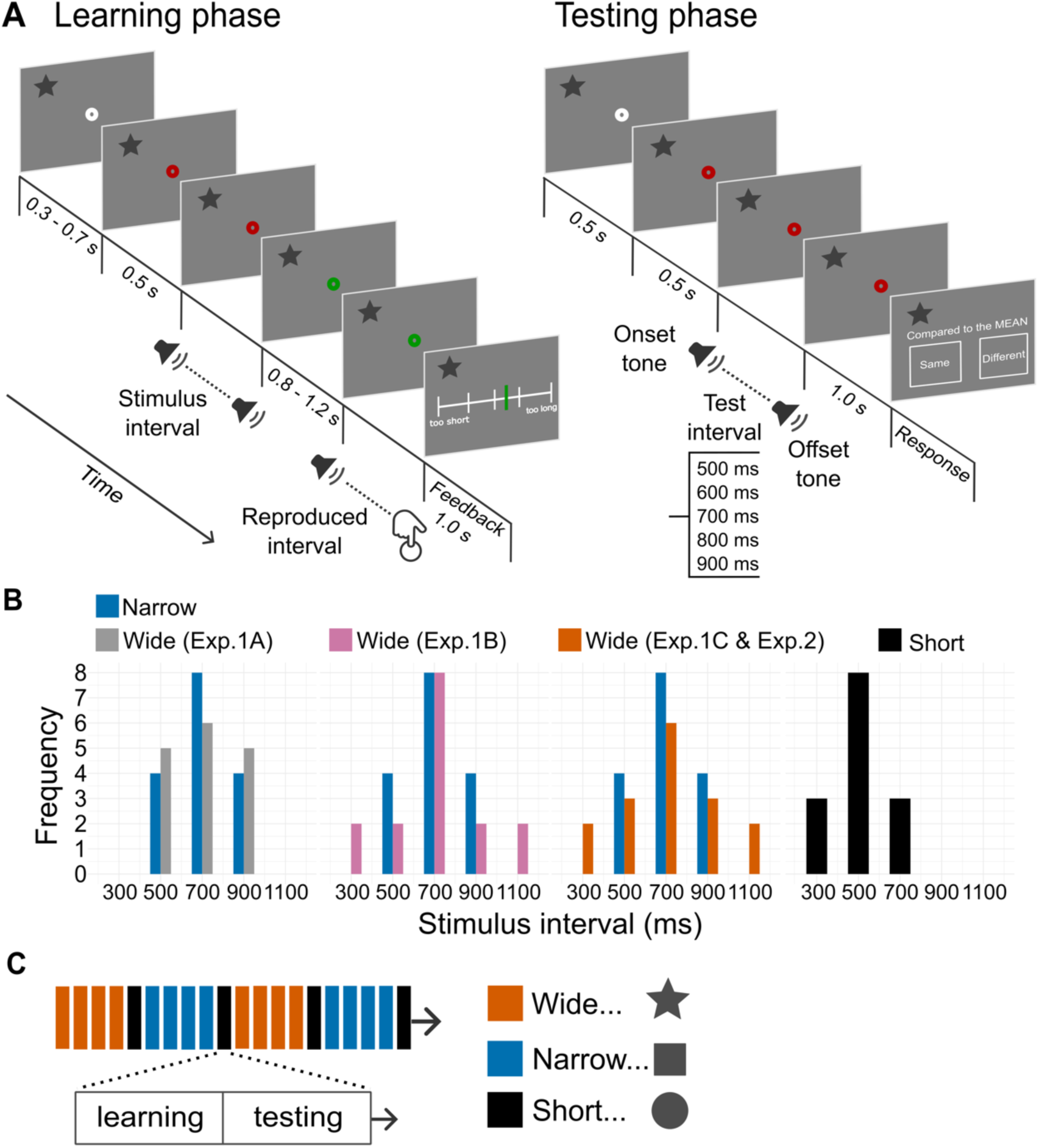
Experimental design and procedure. *(A) Trial structure.* The task consisted of two successive phases: a learning phase and a testing phase. In the learning phase, participants performed an interval reproduction task. They measured a stimulus interval defined by two tones. After the third tone, they were instructed to reproduce the interval by pressing a button to indicate the offset of the reproduced interval, followed by feedback on the accuracy of their reproduction. In the testing phase, participants performed a temporal generalization task where they judged whether a test interval was the “same” or “different” compared to the mean of the stimulus intervals that they had measured in the preceding learning phase. One of the three symbols—a star, a square, or a circle—was displayed on the top left of the screen during the entire task to indicate the stimulus distribution of the current block. *(B) Stimulus distribution.* Each version of experiment (Experiment 1A, 1B, 1C, and 2) comprised the three stimulus distributions: narrow, wide, and short. The narrow (blue) and short (black) distribution was identical across experiments, whereas the wide distribution varied across Experiment 1A (gray), 1B (pink), and 1C (orange). The stimulus distribution of Experiment 2 was identical to that of Experiment 1C. *(C) Experiment structure.* One experimental session consisted of 20 blocks, where each block comprised a learning phase and a testing phase. The colors (orange, blue, and black) indicate the stimulus distribution of the corresponding block. The short distribution was always tied to the circle symbol, while the narrow and wide distributions were tied to either the star or square, with the associations counterbalanced across participants.

#### Procedure

The task consisted of two successive phases: a learning phase and a testing phase.

*The learning phase*. Participants performed an auditory interval reproduction task (see Fig. 1A). Each trial started with a white fixation dot shown for 300 – 700 ms (uniformly sampled). The fixation dot then turned red and remained red before (500 ms) and during the presentation of two 50 ms tones. The two tones indicated the onset and offset of the stimulus interval, defined as the inter-stimulus-interval between the two tones. After the offset tone, the fixation dot turned green for 800 – 1200 ms (uniformly sampled) before the presentation of another 50 ms tone and remained green until the participant’s button press. Participants were instructed to press a button so that the interval between this third tone and their button press—i.e., reproduced interval— matched the earlier presented stimulus interval as closely as possible. Upon giving their responses, participants received feedback for 1 second, indicating the accuracy of their reproduction. The relative error was defined as follows: (*reproduced interval - stimulus interval*) / *stimulus interval* and was displayed as a bar on a continuous scale demarcated by five values: -0.4 (“too short”), -0.15, 0, 0.15, 0.4 (“too long”). The bar was shown in green if the relative error was between -0.15 and 0.15, and in red otherwise.

*The testing phase*. Participants performed a temporal generalization task where they answered whether a given interval was the “same” or “different” compared to the mean of the stimulus intervals that they had encountered in the learning phase (Fig. 1A). Each trial started with a white fixation dot shown for 500 ms, which then turned red and remained for another 500 ms. Following this, two 50 ms tones (the onset tone and the offset tone) were presented to mark the onset and offset of the test interval. After a delay of 1000 ms, empty squares labeled “Same” and “Different” appeared on either side of the screen. Participants were instructed to press the button corresponding to the “Same” side of the gamepad if they thought the test interval was the same as the mean, and the “Different” side if they thought it was different from the mean. The left and right positions of the “Same” and “Different” labels on the screen were counterbalanced across trials.

*Conditions*. Before starting the learning phase, participants saw one of the three symbols—a star, a square, or a circle—displayed in the center of the screen. This indicated that the stimulus intervals in the upcoming learning phase would be drawn from a specific distribution tied to that symbol and that participants would be asked about the mean of that distribution in the following testing phase. The symbol was always displayed on the top left of the screen during the entire task (Fig. 1A). The three symbols, star, square, and circle were tied to the three different stimulus distributions during the learning phase: narrow, wide, and short (Fig. 1B). The narrow and short distributions were identical across Experiments 1A, 1B, and 1C: the narrow distribution consisted of intervals of 500, 700, and 900 ms, presented 4, 8, and 4 times each (mean = 700 ms), while the short distribution consisted of intervals of 300, 500, and 700 ms, presented 3, 8, and 3 times each (mean = 500 ms). In contrast, the wide distribution had the same mean as the narrow distribution but had different variability, which differed across Experiments 1A, 1B, and 1C (Fig. 1B). In Experiment 1A, the wide distribution comprised 500, 700, and 900 ms intervals, presented 5, 6, and 5 times each (mean = 700 ms). In Experiment 1B, it comprised 300, 500, 700, 900, and 1100 ms intervals, presented 2, 2, 8, 2, and 2 times each (mean = 700 ms). In Experiment 1C, it comprised 300, 500, 700, 900, and 1100 ms intervals, presented 2, 3, 6, 3, and 2 times each (mean = 700 ms). Thus, the wide distribution in Experiment 1A had the same width as the narrow distribution but with lower presentation frequency of the mean interval (700 ms) compared to the narrow distribution. Conversely, the wide distribution in Experiment 1B had the same presentation frequency of the mean interval but a greater width compared to the narrow distribution. In Experiment 1C, the wide distribution had a greater width as well as a lower presentation frequency of the mean interval. Therefore, the variance of the distributions gradually increased in the order of narrow, wide (Experiment 1A), wide (Experiment 1B), and wide (Experiment 1C). The stimulus presentation was randomized within each distribution. The short distribution was always tied to the circle symbol, while the narrow and wide distributions were tied to either the star or square, with the associations counterbalanced across participants. While participants were informed that the stimulus distributions were tied to the three symbols, they were not informed about their content (i.e., narrow, wide, and short). The test intervals in the testing phase comprised intervals of 500, 600, 700, 800, and 900 ms, and the presentation frequency, four times each, was constant across all experiments and conditions.

One experimental session consisted of 20 blocks, where each block comprised a learning phase and a testing phase (Fig. 1C). Each block included 16 trials (narrow and wide) or 14 trials (short) for the learning phase and 20 trials for the testing phase (5 test intervals presented 4 times each). The 20 blocks were further grouped into 4 chunks of 5 blocks each. Each chunk started with 4 consecutive blocks of either narrow or wide distribution, followed by one block of short distribution. Chunks containing narrow and wide distributions were alternated; for example, when the first chunk contained narrow, it was followed by a chunk containing wide. Whether the first chunk started with a narrow or wide distribution was counterbalanced across participants. Participants could take self-timed breaks after each testing phase. Before the formal experiment, participants performed a practice block comprising 12 trials of the learning phase (600, 800, and 1000 ms intervals, each presented 2, 8, and 2 times) and 10 trials of the testing phase (600, 700, 800, 900, 1000 ms intervals, each presented 2 times). In total, each test interval had 32 data points for the narrow and wide conditions and 16 data points for the short condition.

The reason for performing four consecutive blocks of the narrow and wide distribution was to solidify the learning of the distributions. Furthermore, including the short distribution before switching between the narrow and wide aimed to make participants “forget” the preceding learning, preventing them from confusing the narrow and wide distributions. Additionally, having a distribution with a different (short) mean interval allowed us to test if participants correctly estimated the mean of different distributions.

#### Data analysis

##### The learning phase (reproduction task)

Trials with reproduced intervals shorter than 120 ms (40% shorter than the shortest interval) and longer than 1540 ms (40% longer than the longest interval) were excluded from the analysis (0.7, 1.2, and 1.5 % of the data of Experiments 1A, 1B, and 1C, respectively). To test whether reproductions were influenced by the distribution (narrow and wide), we fitted linear mixed-effect models (LMMs) using the *lme4* package (Bates et al., 2015) in R version 4.2.3 (R Core Team, 2023), including ‘stimulus interval’, ‘distribution’, and their interaction term as fixed factors. To improve the model fit, ‘stimulus interval’ was centered at 700 ms, and ‘distribution’ was recoded using effect coding (-0.5 for narrow and 0.5 for wide). The significant fixed factors we report below are based on the models that include a random slope of ‘participant’ for the corresponding fixed factors in addition to a random intercept.

##### The testing phase (generalization task)

Trials with reaction times (RTs) shorter than 150 ms and longer than 4000 ms were excluded (0.7, 1.0, and 2.4 % of the data of Experiments 1A, 1B, and 1C, respectively). To assess whether the distribution (narrow and wide) affected the “same”/“different” responses, we fitted generalized linear mixed-effects models (GLMMs) using *glmmTMB* package (Brooks et al., 2017) with the binomial family and a logit link function. Participants’ responses were coded as 1 when they answered “same” and 0 when they answered “different”. Since the probability of the “same” responses was expected to peak at the true mean (700 ms), test intervals were recoded as ‘absolute difference’ (0 for 700 ms, 1 for 600 and 800 ms, and 2 for 500 and 900 ms) and ‘shorter longer’ (-0.5 for 500 and 600 ms, 0 for 700 ms, and 0.5 for 800 and 900 ms). Thus, GLMMs predicting response (1 and 0) included ‘absolute difference’, ‘shorter longer’, ‘distribution’ (-0.5 for narrow and 0.5 for wide), and their interaction terms as fixed effect factors. In addition to a random intercept of ‘participant’, we sequentially added random slopes for the fixed factors that were significant in simpler models and tested whether they improved the model with a likelihood ratio test. The significant fixed factors we report below are those from the best-fitting models, which include a random slope for the corresponding fixed factors.

### Experiment 2 (EEG)

#### Participants

Twenty-three first-year psychology students, who did not participate in Experiment 1, participated for partial course credit. The sample size was based on previous EEG experiments from our group (Damsma et al., 2021; Kononowicz & van Rijn, 2014). One participant was excluded from the analysis due to excessive artifacts in the EEG data, resulting in the final sample of 22 participants (mean age = 21.4 years; 16 female). All participants reported normal or corrected-to-normal vision, normal hearing, and no history of psychiatric or neurological conditions. The experiment was approved by the Ethical Committee of the Faculty of Behavioural and Social Sciences (PSY-2324-S-0091). Written informed consent was obtained before the experiment.

#### Apparatus, stimuli, and procedure

The experimental setup was identical to Experiment 1 except for the recording of EEG throughout the experiment. The stimulus distribution was identical to that of Experiment 1C: the wide distribution had a greater width and a lower presentation frequency of the mean interval compared to the narrow distribution.

#### EEG acquisition and preprocessing

EEG data were recorded with TMSi Refa 8-64 amplifier using 64-channel Ag/AgCl electrodes placed following the international 10-20 system (WaveGuard EEG cap, eemagine Medical Imaging Solutions) at the sampling rate of 1000 Hz. AFz and CPz served as a ground and an online reference electrode, respectively. The electrooculogram was recorded from an electrode on the bottom of the left eye. The impedance of all electrodes was kept below 10 kΩ throughout the experiment.

Preprocessing was performed using FieldTrip toolbox (Oostenveld et al., 2010) in MATLAB. Epochs were defined as segments from -500 to 1500 ms relative to the presentation of the onset and offset tones of the test intervals during the testing phase (the onset epochs and the offset epochs). The data were re-referenced to the average of the left and right mastoids and filtered by a 0.01-80 Hz band-pass Butterworth filter. Epochs that contained signals exceeding an amplitude range of 300 μV and participants with more than 50 % of such epochs were excluded from further analysis (8.7 and 7.5 % of the data were discarded for the onset and offset epochs, respectively, and one participant was excluded from the further analysis). Ocular artifacts were corrected using independent component analysis based on typical scalp topography and time course.

#### Data analysis

##### Behavioral analysis

The behavioral data analysis was identical to that of Experiment 1. We excluded 0.5 and 1.7 % of the data from the learning and testing phases, respectively.

##### ERP analysis

All ERP analyses we will report here are focused on the testing phase. Based on previous research (Damsma et al., 2021; Kononowicz & van Rijn, 2014), we focused all ERP analyses on the front-central electrodes (Cz, C1, C2, FCz, FC1, FC2). All ERPs were calculated on a single-trial basis and were averaged across participants, test intervals, and distribution (narrow and wide) when appropriate.

ERPs locked to the interval onset (onset-tone locked ERPs) were filtered by a 10 Hz Butterworth low-pass filter and corrected to the average amplitude in the 100 ms baseline window before the interval onset (Damsma et al., 2021; Kononowicz & van Rijn, 2014). CNV amplitude was determined by calculating the mean amplitude over a window from 50 to 0 ms before the interval offset (i.e., the offset tone) for each test interval, based on an earlier finding that CNV immediately before stimulus onset reflects the state of preparation (Boehm et al., 2014). ERPs locked to the interval offset (offset-tone locked ERPs) were filtered by a 1-20 Hz band-pass Butterworth filter to attenuate CNV-based contamination and corrected to the average amplitude in the baseline window from -50 to 50 ms relative to the interval offset (Damsma et al., 2021; Kononowicz & van Rijn, 2014). Offset-tone locked N1, offset-tone locked P2, and LPC amplitude were defined as the mean amplitude between 70 and 160 ms, 140 and 300 ms, and 300 and 500 ms after the interval offset, respectively. The choice of these time windows of interest was based on previous studies (Baykan et al., 2023; Kononowicz & van Rijn, 2014; Ofir & Landau, 2022) and the visual inspection of the averaged waveform across all trials.

To test whether ERP amplitudes were influenced by the test interval and distribution, we fitted LMMs similar to those used in behavioral analysis, predicting single-trial ERP amplitude based on ‘test interval’ (centered at 700 ms), ‘distribution’ (-0.5 for narrow and 0.5 for wide), and their interaction term as fixed factors. Additionally, to examine a potential non-linearity in the effect of test interval, we fitted alternative models that included ‘absolute difference’ (0 for 700 ms, 1 for 600 and 800 ms, and 2 for 500 and 900 ms), ‘shorter longer’ (-0.5 for 500 and 600 ms, 0 for 700 ms, and 0.5 for 800 and 900 ms), ‘distribution’, and their interaction terms as fixed effect factors. We first compared these two types of models for each ERP component using a likelihood ratio test. Then, for the winning model, we sequentially added a random slope of ‘participant’ for the corresponding fixed factors to test whether they improved the model. We report the significant fixed factors from the best-fitting models, which include a corresponding random slope in addition to a random intercept.

##### Relation between ERP and behavior

We investigated the link between ERPs and behavioral measures based on two approaches: the single-trial analysis using (G)LMM and the individual differences.

##### Single-trial ERP and behavior

To test whether ERP amplitude predicts single-trial response (“same”/“different”), we fitted GLMMs separately for CNV, offset-tone locked N1, offset-tone locked P2, and LPC. These models extended the best-fitting models from the behavioral analysis by incorporating the fixed factor ‘ERP amplitude’ and its corresponding random slope. Thus, the models included ‘absolute difference’, ‘shorter longer’, ‘distribution’, ‘ERP amplitude’, and their interaction terms as fixed factors.

Additionally, to test whether ERP amplitude predicts single-trial reaction time (RT), we fitted LMMs. These models included ‘absolute difference’, ‘shorter longer’, ‘ERP amplitude’, and ‘response’ (-0.5 for “same” and 0.5 for “different”), and their interaction terms as fixed factors but not the term ‘distribution’, based on preliminary analyses of the RT data (Appendix Figure A1). Before analysis, RTs were log-transformed, and ERP amplitudes were z-transformed per participant. All models included a random intercept of ‘participant’. The significant fixed factors we report are based on the models that include corresponding random slopes of ‘participant’.

##### Individual differences

Is there a relation between how much individuals’ performance is affected by the narrow versus wide distribution and their between-condition ERP difference? Results of Experiment 1 suggested that the behavioral difference between the narrow and wide conditions was most evident at the 700 ms (the mean) and 500 ms (the shortest) test intervals. Based on this observation, we defined “the behavioral distribution effect” for each participant, by computing the difference in the proportion of “same” responses between 700 ms and 500 ms (i.e., the “height”) for each condition (i.e., narrow and wide) and calculating its difference between conditions (narrow - wide). Similarly, we defined “the EEG distribution effect” by computing the across-trial mean ERP amplitude aggregated across test intervals for each condition and calculating its difference between conditions (wide - narrow). We tested an interindividual link between the behavioral distribution effect and the EEG distribution effect in the CNV, offset-tone locked N1, offset-tone locked P2, and LPC using a two-tailed Pearson’s correlation.

## RESULTS

### Experiment 1

#### Learning phase (reproduction task)

Figure 2A illustrates the mean reproduced interval across participants. The results of the LMM showed that the reproduced interval increased with the stimulus interval in all versions of Experiment 1 (*β* = 0.696, *t* = 16.725, *p* <.001, for 1A; *β* = 0.656, *t* = 16.343, *p* < .001, for 1B; *β* = 0.763, *t* = 31.849, *p* < .001, for 1C). The interaction between stimulus interval and distribution was not significant in Experiments 1A and 1B (*β* = -0.020, *t* = -0.983, *p* = 0.326 and *β* = -0.036, *t* = -1.741, *p* = .081, respectively), but significant in Experiment 1C (*β* = -0.076, *t* = - 2.773, *p* = .009), indicating that the slope was lower (i.e., the central tendency was smaller) for the wide distribution compared to the narrow distribution in Experiment 1C.

**Figure 2.**
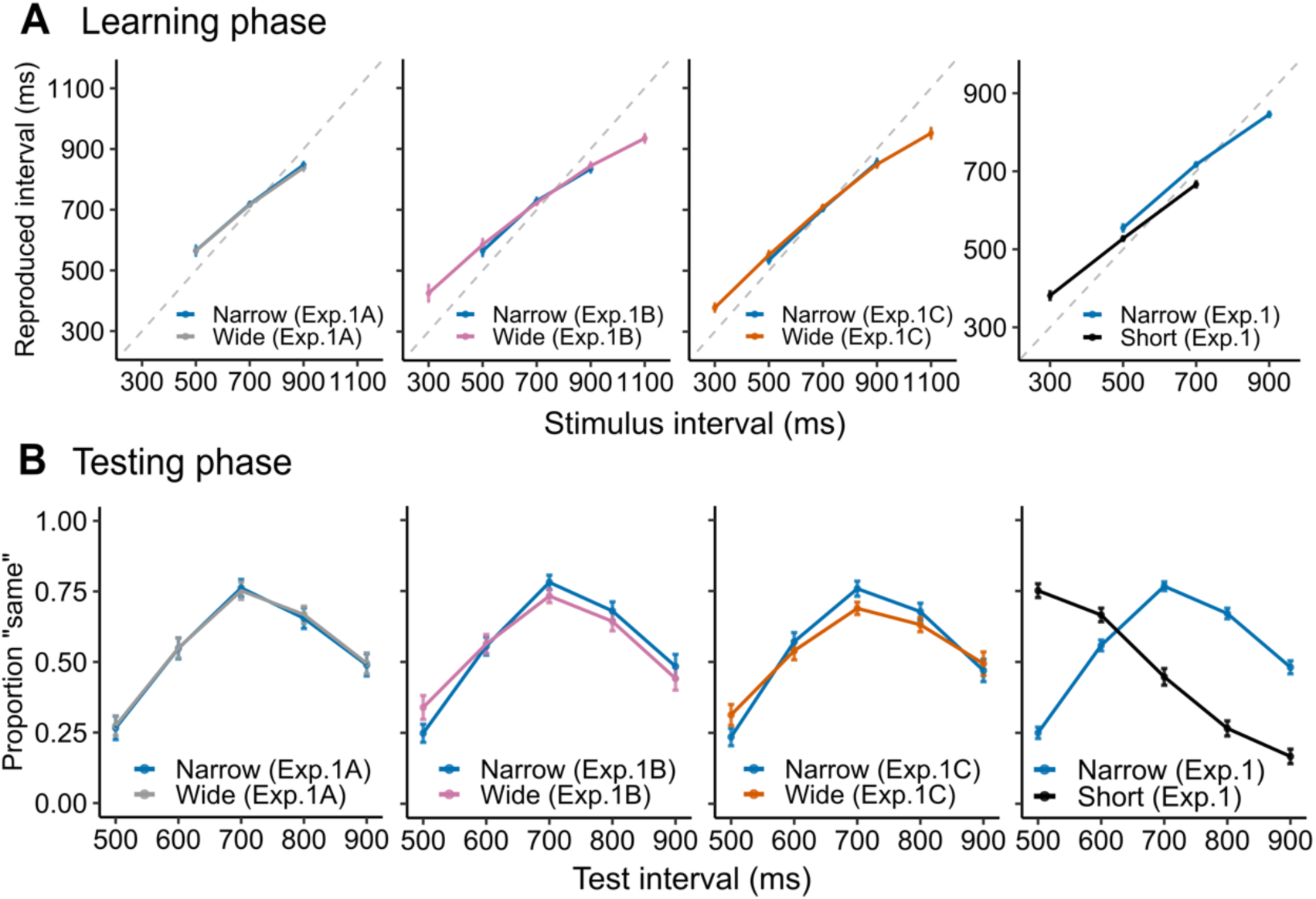
Results of Experiment 1. (A. Learning Phase) Across-participant mean reproduced intervals in the learning phase (interval reproduction task) for each experiment, color-coded by distribution. (B. Testing Phase) Across-participant mean proportions of “same” responses in the testing phase (temporal generalization task) for each experiment, color-coded by distribution. For the narrow and short distribution, reproduced intervals and proportions of “same” responses are averaged across Experiments 1A, 1B, and 1C and plotted in the top-right and bottom-right panels, respectively. Error bars indicate SEM.

#### Testing phase (generalization task)

Figure 2B illustrates the mean proportion of “same” responses across participants. Overall, the data of the narrow and wide conditions showed an inverted V-shape peaking at the 700 ms test interval, with a noticeable asymmetry with higher proportions on the positive side. The results of the GLMM confirmed this pattern. In all versions of Experiment 1 (1A, 1B & 1C), the probability of “same” responses peaked at 700 ms and decreased as the test interval deviated from 700 ms (main effect of absolute difference: *β* = -1.035, *z* = -8.776, *p* < .001, for 1A; *β* = -1.009, *z* = -9.410 *p* < .001, for 1B; *β* = -0.895, *z* = -9.677 *p* < .001, for 1C). Moreover, the probability of “same” responses was higher for the longer test intervals compared to shorter ones (interaction between absolute difference and shorter longer: *β* = 0.742, *z* = 3.685, *p* < .001, for 1A; *β* = 0.452, *z* = 2.799, *p* = .005, for 1B; *β* = 0.668, *z* = 3.881, *p* < .001, for 1C). These results suggest that participants accurately estimated the distributions’ mean while also showing a tendency to select longer intervals as the mean more often than shorter ones.

The effect of distribution varied between the experiments. In Experiment 1A, there was no significant difference between the narrow and wide distributions (main effect of distribution: *β* = -0.028, *z* = -0.307, *p* = .758). In contrast, in Experiments 1B and 1C, the probability of “same” responses was lower for the wide distribution compared to the narrow distribution (main effect of distribution: *β* = -0.298, *z* = -1.988, *p* = .046 and *β* = -0.569, *z* = -3.787, *p* < .001, for 1B and 1C, respectively). Furthermore, the effect of distribution interacted significantly with that of test interval (interaction between absolute difference and distribution: *β* = 0.210, *z* = 2.034, *p* = .041 and *β* = 0.421, *z* = 4.289, *p* < .001, for 1B and 1C, respectively), confirming the pattern that the proportion of “same” responses was lower at the 700 ms test interval and higher at the 500 ms test interval in the wide condition compared to the narrow condition (Fig. 1B, Experiments 1B and 1C). Moreover, the data of the short distribution showed a peak at the test interval of 500 ms and decreased with the longer test intervals (Fig. 2B), suggesting that participants accurately estimated the mean of the short distribution. Taken together, these results indicated that participants were able to accurately estimate the mean of the distributions. Furthermore, although the actual mean remained constant, the variability of the estimated mean increased with the variability of the distribution.

### Experiment 2

#### Behavior

The experimental setup was identical to that of Experiment 1C. Figure 3A shows the mean reproduced interval across participants during the learning phase. The LMM showed that the reproduced interval increased with the stimulus interval (*β* = 0.735, *t* = 24.471, *p* < .001). However, unlike in Experiment 1C, the interaction between stimulus interval and distribution was not significant (*β* = 0.019, *t* = 0.995, *p* = .320), possibly due to the smaller sample size compared to Experiment 1C. Nevertheless, the reproduction data from Experiment 2 appear qualitatively similar to that from Experiment 1C (Fig. 3A). Figure 3B illustrates the mean proportion of “same” responses across participants during the testing phase. Consistent with Experiment 1C, the GLMM revealed the significant main effect of absolute difference (*β* = -0.924, *z* = - 7.663, *p* < .001), the interaction between absolute difference and shorter longer (*β* = 0.805, *z* = 4.272, *p* < .001), the main effect of distribution (*β* = -0.454, *z* = -2.163, *p* = .030), and the interaction between absolute difference and distribution (*β* = 0.335, *z* = 2.561, *p* = .010). Thus, the behavioral results of the testing phase in Experiment 2 replicated the main findings from Experiment 1C: the variability of the estimated mean increases as a function of the variability of the distribution.

**Figure 3.**
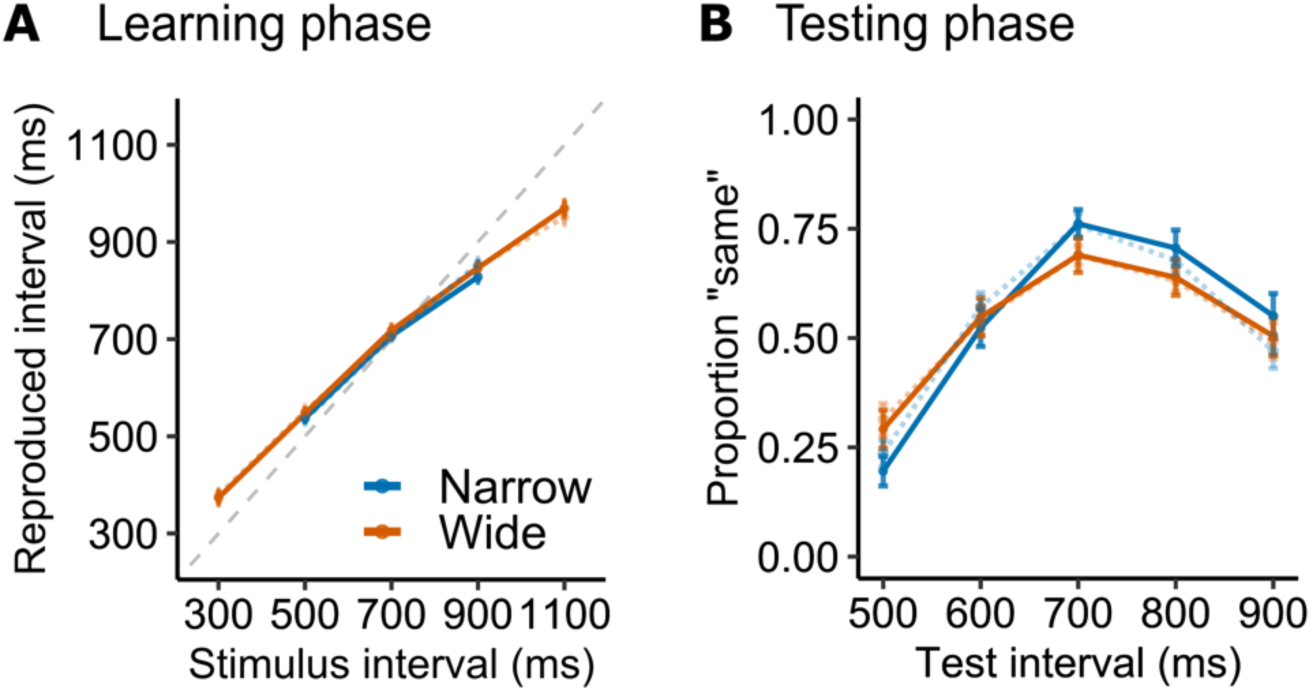
Behavioral results of Experiment 2. (A. Learning Phase) Across-participant mean reproduced intervals in the learning phase. (B. Testing Phase) Across-participant mean proportions of “same” responses in the testing phase. The colors indicate the stimulus distribution (blue, narrow; orange, wide). The data from Experiment 2 are shown with the solid lines, while the data from Experiment 1C are overlaid with the dotted lines for comparison. Error bars indicate SEM.

#### ERPs

Figures 4A and 4B show the average onset-tone-locked ERP and offset-tone-locked ERP in the testing phase, separately for the narrow and wide conditions. The LMM results did not reveal a significant effect of distribution (i.e., narrow and wide) for any ERP components. Nevertheless, we present the data separately for distribution in Figure 4 for the sake of transparency and to allow for comparisons with the behavioral data. The across-participant mean amplitudes of the CNV, offset-tone locked N1, offset-tone locked P2, and LPC are shown in Figure 4C. The LMM results indicated that neither a main effect nor an interaction was significant for the CNV and offset-tone locked N1. However, the offset-tone locked P2 increased proportionally with the test interval (main effect of test interval: *β* = 0.361, *t* = 3.776, *p* = .001). The LPC was lowest at the test interval of 700 ms and tended to plateau for the longer test intervals (main effect of absolute difference: *β* = 0.388, *t* = 2.395, *p* = .026; interaction between absolute difference and shorter longer: *β* = -1.362, *t* = -4.095, *p* < .001).

**Figure 4.**
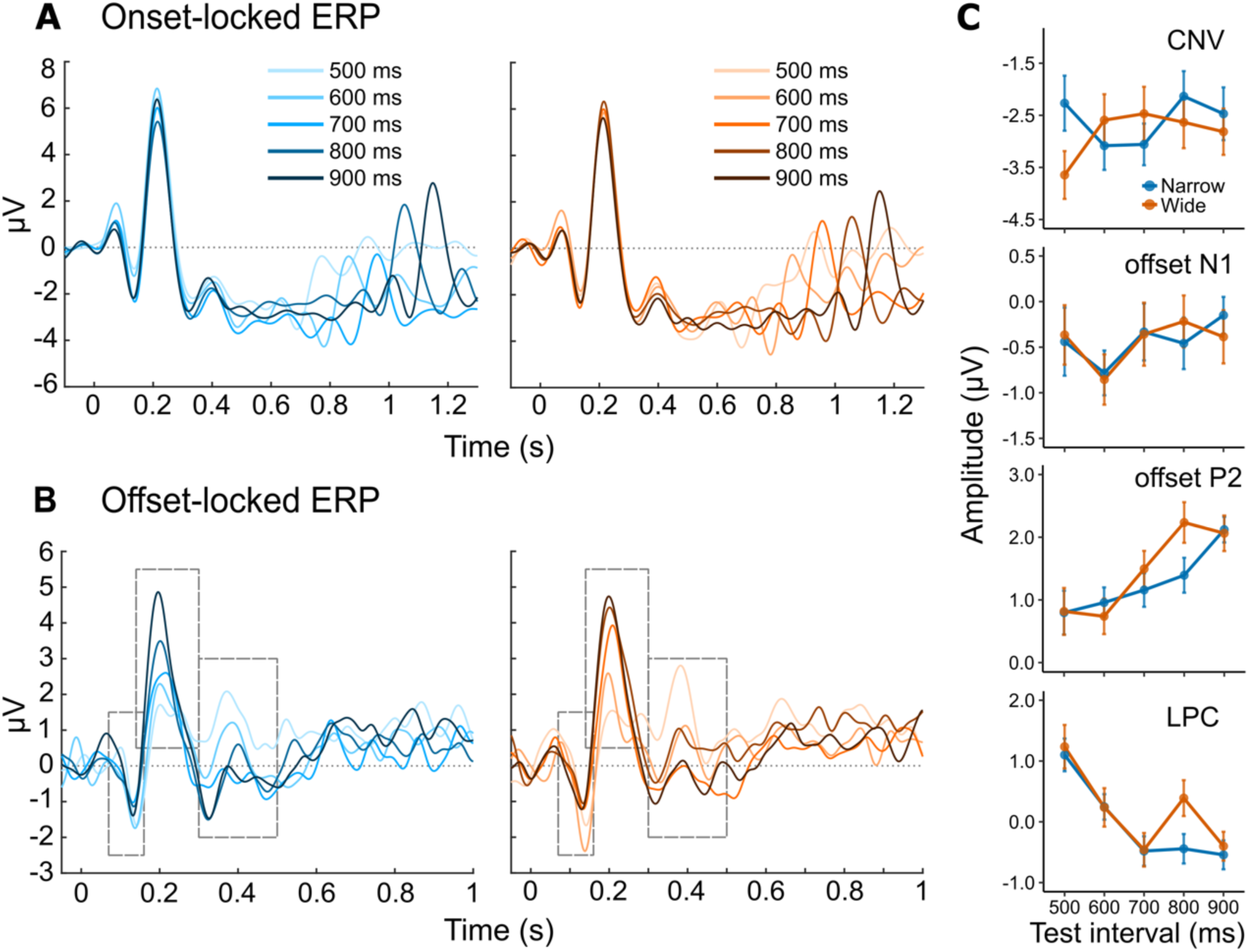
Across-participant mean ERPs at the front-central electrodes (Cz, C1, C2, FCz, FC1, FC2) in the testing phase. (A) Onset-tone locked ERP; (B) Offset-tone locked ERP. The left (blue) and right (orange) panels show the waveforms in the narrow and wide distribution conditions, respectively, for the different test intervals. The gray empty squares in the bottom panels represent the analysis time windows for the offset-tone locked N1, offset-tone locked P2, and LPC. (C) Across-participant mean amplitudes of CNV, offset-tone locked N1, offset-tone locked P2, and LPC as a function of test interval and distribution (blue, narrow; orange, wide). Error bars indicate SEM.

#### Relation between ERP and behavior

*Single-trial ERP and behavior.* The GLMM results indicated that the single-trial offset-tone locked P2 predicted the participants’ responses (“same”/“different”), with a higher P2 amplitude increasing the probability of a “different” response (main effect of P2 amplitude: *β* = -0.165, *z* = -2.644, *p* = .008). Similarly, a lower (i.e., more positive) N1 amplitude was associated with a higher probability of a “different” response (main effect of N1 amplitude: *β* = -0.176, *z* = -2.907, *p* = .003), although this effect interacted with test interval (interaction between absolute difference and N1 amplitude: *β* = 0.132, *z* = 2.798, *p* = .005). No evidence that the CNV and LPC predicted the responses was found. To illustrate the findings, we plotted the across-participant mean amplitudes of the offset-tone locked N1 and P2, separately for trials with the “same” and “different” responses in Figure 5A. Notably, the P2 amplitude showed a clear increase with the test interval in trials where participants responded “different”, while this increase seemed less clear in trials where they responded “same”. To verify this pattern, we performed a post-hoc analysis by fitting an LMM predicting P2 amplitude with test interval, response (-0.5 for “same” and 0.5 for “different”), and their interaction as fixed factors, along with a random intercept and random slopes. The results confirmed the visual inspection, revealing a significant interaction between test interval and response (*β* = 0.322, *t* = 2.279, *p* = .027) in addition to a significant main effect of interval (*β* = 0.382, *t* = 3.950, *p* < .001) and response (*β* = 0.470, *t* = 2.201, *p* = .033). A similar pattern was not observed for the N1.

**Figure 5.**
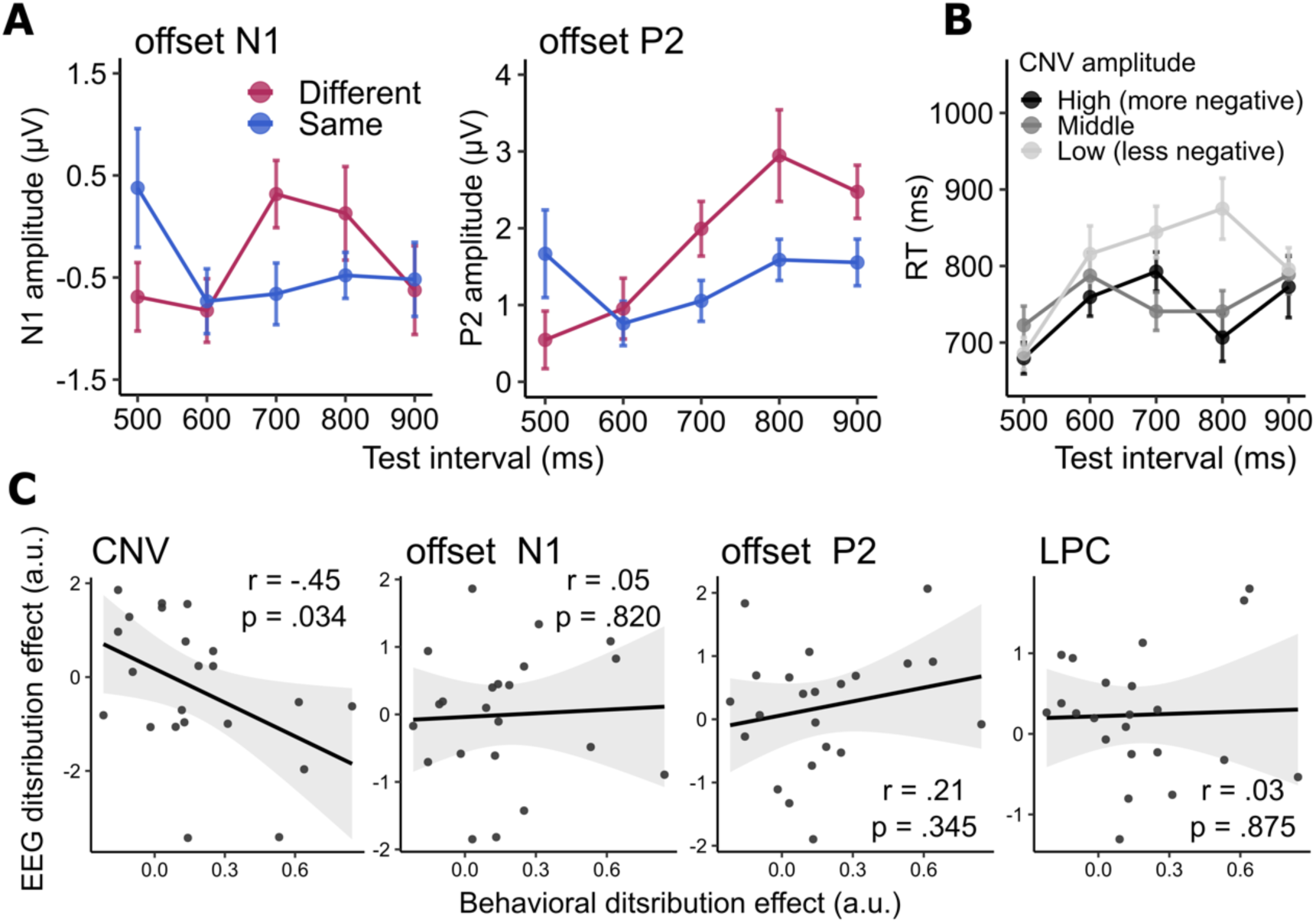
Relation between ERP and behavior. (A) Offset-tone locked N1/P2 and response (“same” and “different”). The left and right panels show across-participant mean amplitudes for the offset-tone locked N1 and P2, respectively, as a function of test interval and response (magenta, “different”; blue, “same”). (B) CNV and RT. Across-participant mean RTs are plotted as a function of test interval and CNV amplitude. Trials were divided into tertiles based on the CNV amplitude (high, middle, and low) per participant and test interval. Error bars indicate SEM. (C) Correlation between participants’ behavioral distribution effect and their EEG distribution effect in the CNV, offset-tone locked N1, offset-tone locked P2, and LPC. The solid lines indicate the best-fitting lines and the shaded regions the 95% confidence intervals. Dots represent individual participants. Pearson’s r and associated p-values are reported.

Additionally, we tested whether the single-trial ERP predicted RT on each trial. Figure 5B illustrates the relationship between CNV and RT. For visualization purposes, all trials were divided into tertiles based on CNV amplitude (high, middle, and low) per participant and test interval, and the mean RT was plotted. The LMM results revealed that the single-trial CNV was predictive of the RT, with a higher (i.e., more negative) CNV amplitude associated with a faster RT (main effect of CNV amplitude: *β* = 0.068, *t* = 4.055, *p* < .001). Furthermore, this effect interacted with the test interval, with the effect evident when the test interval was close to 700 ms (interaction between absolute difference and CNV amplitude: *β* = -0.024, *t* = -2.244, *p* = .027). The LMM showed no evidence that the offset-tone locked N1, P2, and LPC predicted RTs.

*Individual differences.* Figure 5C shows the correlation between “the behavioral distribution effect” and “the EEG distribution effect” in the CNV, offset-tone locked N1, offset-tone locked P2, and LPC. The two-tailed Pearson’s correlation coefficient indicated that these measures were correlated for the CNV (*r* = -0.45, *t*(20) = -2.263, *p* = .034), suggesting that individuals whose performance was more affected by the distributions exhibited higher (i.e., more negative) CNV amplitudes in the wide condition relative to the narrow condition. No significant correlation was found for the offset-tone locked N1 (*r* = 0.05, *t*(20) = 0.230, *p* = .820), P2 (*r* = 0.21, *t*(20) = 0.965, *p* = .345), and LPC (*r* = 0.03, *t*(20) = 0.158, *p* = .875).

## DISCUSSION

Previous work on temporal context effects has shown that humans are highly sensitive to temporal regularities. Temporal perception, for instance, is influenced by the underlying stimulus distributions (Duhamel et al., 2023; Jazayeri & Shadlen, 2010; Miyazaki et al., 2005; Ryan, 2011) and the history of presented stimuli (Glasauer & Shi, 2021, 2022; Rhodes et al., 2023; Taatgen & van Rijn, 2011). Particularly, Acerbi et al. (2012) suggested that humans can learn up to third-order statistics (i.e., the mean, variance, and skewness) of temporal distributions. In typical experiments, participants are required to time stimuli on each trial, revealing a dynamic adaptation process in which the current temporal estimate is modulated on a trial-by-trial basis (de Jong et al., 2021; Glasauer & Shi, 2021; Zimmermann & Cicchini, 2020). However, in real-world scenarios, learning temporal regularities does not always occur as a dynamic adaptation to samples; it often involves extracting and retaining summary statistics, such as the mean, of a distribution to guide future behavior. In this study, we focus on such learning processes by explicitly asking participants to extract the mean of distributions of time intervals. Additionally, we examine whether the variability of these distributions affects how the mean is represented.

In the experiments, participants first reproduced time intervals drawn from one of two distributions that had the same mean but different variability (i.e., one narrow and one wide distribution). They then compared the mean of the distribution to subsequently presented test intervals in a temporal generalization task. The temporal generalization data showed a peak at the actual mean interval, a pattern typically observed in temporal generalization tasks that use only one standard duration (Bannier et al., 2019; Droit-Volet & Coull, 2016; Özoğlu & Thomaschke, 2023; Wearden, 1992), indicating that participants accurately estimated the mean of the temporal distributions. Crucially, these estimates of the mean were influenced by the distribution’s variability operationalized as the width and the presentation frequency of the intervals. In Experiment 1A, where the two distributions differed only in the presentation frequency of the mean interval, we failed to find a distribution effect on the estimates. However, in Experiments 1B and 1C, where the distributions differed in width, or in both width and presentation frequency, the variability did affect the estimates of the mean. These findings indicate that participants’ subjective estimates of the mean were sensitive to the distribution’s variability—particularly its width—even when the variability was irrelevant to the mean itself. Notably, higher sensitivity to the width relative to the presentation frequency suggests that the summary representation of the mean of time intervals is more strongly influenced by the tail of the distribution than by the height of the peak. These results align with previous findings that the central tendency effects become more pronounced as the width of the stimulus distribution increases (Duhamel et al., 2023; Miyazaki et al., 2005), implying that summary representations for time intervals and temporal context effects may share common computational mechanisms. As we will discuss later, these results support the possibility of a unified description within a Bayesian framework.

While previous studies have also demonstrated that humans can extract the mean of multiple time intervals (Curtis & Rule, 1977; Ren et al., 2020; Schweickert et al., 2014; Wearden & Jones, 2007), our work provides the first evidence that the variability of these estimates increases with the variability of the distribution of time intervals. In the earlier study by Wearden & Jones (2007), participants were also asked to estimate the mean of three sample durations, which varied in their spacing (i.e., variability) across conditions. In their experiment, three sample durations were each presented once at the beginning of a block, after which participants performed a temporal generalization task, judging whether the test durations matched the mean of the three sample durations. The distance between the sample durations varied across conditions, ranging from 25 ms to 450 ms. Some conditions included the mean within the sample durations (i.e., the middle sample duration was equidistant from the others) while others did not (i.e., the spacing between the sample durations was random). Interestingly, while their participants indeed correctly extracted the mean, they did not seem to be influenced by the variability of the sample durations in any of the conditions.

It is important to note, however, that the purpose of Wearden and Jones (2007) study was to explore the nature of humans’ internal time scale by examining whether participants’ estimates of the mean corresponded to the arithmetic mean or not, whereas our study aimed to investigate how humans learn temporal distributions by testing whether estimates of the mean are influenced by the distribution’s variability. This distinction led to key differences in experimental design: while Wearden & Jones (2007) presented three different sample durations only once, we presented five different sample durations (with 100 ms distance between samples) multiple times, with a total of 16 samples. In other words, while presenting three sample durations was sufficient for their objective, we needed to present more samples to create a robust temporal distribution within which the effects of variability could be tested. It is therefore likely that the increased number of samples and presentation repetition in our experiment resulted in greater uncertainty in the representation of the mean interval, making it more sensitive to the variability of the samples. Additionally, the patterns in our temporal generalization data are distinct from those previously observed where memory load was increased by requiring participants to remember two different standards (Jones & Wearden, 2004) or where response thresholds were altered by changing the spacing of test intervals (Ferrara et al., 1997; Uraguchi et al., 2022). The present data are therefore consistent with the interpretation that increased variability in the distribution leads to greater uncertainty in the representation of the mean.

What is the significance of being sensitive to the variability (i.e., variance) when estimating the mean of temporal distributions? A key advantage of retaining information about samples’ variability is that it reflects the uncertainty about the estimated mean (Whitney & Yamanashi Leib, 2018). For example, even if two baseball pitchers have the same average pitch speed, a batter must adjust their preparation differently if the variability around that average differs. The fact that the estimated mean in our study was influenced by the variability suggests that humans can retain the uncertainty about their representation of the mean. This finding provides insights into the models of how humans represent temporal regularities of events. Temporal context effects are often consistent with Bayesian integration models, where current sensory inputs (the likelihood function) are integrated with a prior distribution representing the temporal regularities. Crucially, this integration is weighted according to the relative uncertainties of the likelihood function and the prior (Knill & Pouget, 2004). However, there is still no consensus on how an internal prior distribution should be modeled, or whether individual differences exist in the uncertainty associated with the prior (Glasauer & Shi, 2022; Maaß et al., 2021; Wang et al., 2023). Future research could explore whether the summary representation of the mean of temporal distributions is related to the prior distribution in the Bayesian integration models. Additionally, individual differences in the degree to which the estimated mean is influenced by the distribution’s variability may index individual differences in the uncertainty about the prior expectations.

Using the same behavioral paradigm, we also investigated how learning temporal distributions modulates EEG signals during subsequent temporal judgments (i.e. while estimating the mean). Before turning to the relationship between EEG and behavioral distribution effects, we first addressed the predictive value of the ERPs for single-trial behavior. Our analysis revealed that single-trial offset-tone locked N1 and P2 components predicted participants’ responses. Notably, while P2 amplitude remained relatively constant in trials where the test interval was judged to be the “same” as the subjective mean, it increased in proportion to the actual test interval in trials where the test interval was judged to be “different”. This pattern aligns with the idea that offset P2 indexes the perceived time of the current stimulus (Damsma et al., 2021; Kononowicz & van Rijn, 2014; Kruijne et al., 2021). In contrast, single-trial CNV predicted the RT, with a higher (i.e., more negative) CNV amplitude associated with a faster response on that trial. This finding collaborates with an earlier finding using a perceptual decision-making task (Boehm et al., 2014), supporting the view that CNV reflects the preparation and anticipation for an upcoming event and action (Damsma et al., 2021; Leuthold et al., 2004; Mento, 2013; Scheibe et al., 2009, 2010).

When examining the relationship between EEG signals and behavioral distribution effects, we found that the CNV amplitude calculated from the same time window that predicts RTs correlated with the extent to which an individual’s performance was influenced by the narrow versus wide distribution. Specifically, individuals with a larger behavioral distribution effect exhibited a higher CNV amplitude in the wide distribution condition. Since participants with a larger behavioral distribution effect likely had greater uncertainty about the mean of the wide distribution, it is plausible that these participants tended to expect shorter intervals to represent the mean for the wide distribution. Such an expectation could lead to faster CNV development, resulting in a larger overall amplitude (Damsma et al., 2021). This is consistent with previous findings where faster CNV development was observed when shorter intervals were expected (Breska & Deouell, 2017; Li et al., 2017; Praamstra et al., 2006; Van Der Lubbe et al., 2004). Another possible explanation is that these individuals allocated more attentional resources in the wide condition due to a more uncertain representation of the mean. This interpretation aligns with earlier findings that CNV correlated with selective attention (Wilkinson & Lee, 1972), sustained attention (Thillay et al., 2015), and attention to the temporal aspects of stimuli (Liu et al., 2013), with higher CNV amplitudes indicating a greater attentional allocation.

In contrast to CNV and offset P2, the roles of the offset N1 and LPC were less clear in the current study. Previous work suggests that offset N1 predicts the perceived duration of a stimulus (Kononowicz & van Rijn, 2014) while the LPC reflects temporal decision-making (Bannier et al., 2019; Baykan et al., 2023; Ofir & Landau, 2022; Özoğlu & Thomaschke, 2023). Based on this, we hypothesized that both the offset N1 and LPC would predict participants’ responses. While the offset N1 indeed predict single-trial responses, the observed pattern was less clear compared to that of P2, underscoring the stronger predictive quality of P2 in interval timing (Damsma et al., 2021; Kruijne et al., 2021). As for the LPC, the notable pattern of decreasing amplitude that eventually plateaued as a function of the time interval is consistent with previous findings (Bannier et al., 2019; Baykan et al., 2023; Ofir & Landau, 2022; Özoğlu & Thomaschke, 2023). However, unlike earlier studies, we did not observe a clear link between LPC and behavioral performance. The reason for this discrepancy —whether due to differences in the task design or modality— remains an open question for future investigation.

Finally, future studies are needed to clarify the computational mechanisms through which the mean of time intervals is represented. Unlike the fast and automatic “ensemble perception” of visual features (Whitney & Yamanashi Leib, 2018), extracting the mean of sequentially presented time intervals likely requires cognitive resources such as attention and working memory (Taatgen et al., 2007). It is possible, therefore, that the entire stimulus distribution is initially represented as an internal prior, as is commonly assumed in Bayesian integration models (Glasauer & Shi, 2022; Jazayeri & Shadlen, 2010; Maaß et al., 2021; Petzschner et al., 2015; Wang et al., 2023), with the mean of the prior distribution being estimated through a working memory-based strategy. Further research is needed to develop a unified Bayesian model that can account for both the learning of the distribution and the subsequent extraction of its mean. Alternatively, the mean interval might be intuitively extracted from the stimulus distribution without explicitly representing the entire distribution as a prior (Chen et al., 2018; Ren et al., 2020), akin to ensemble perception of sequentially presented auditory tones (Piazza et al., 2013). Our current data are not sufficient to distinguish between these alternative hypotheses. Nevertheless, our study highlights a new aspect of how humans learn statistical regularities in the environment— specifically, the estimation of the mean of temporal distributions—and provides both behavioral and neural correlates of the summary representations for time intervals.

Bringing together the behavioral and EEG results, our work demonstrates not only that humans can accurately estimate the mean of a temporal distribution, but also that their representation of the mean becomes more uncertain as the variability of the distribution increases, which is neurally reflected through the preparation-related CNV observed during temporal judgments.

## APPENDIX

**Figure A1.**
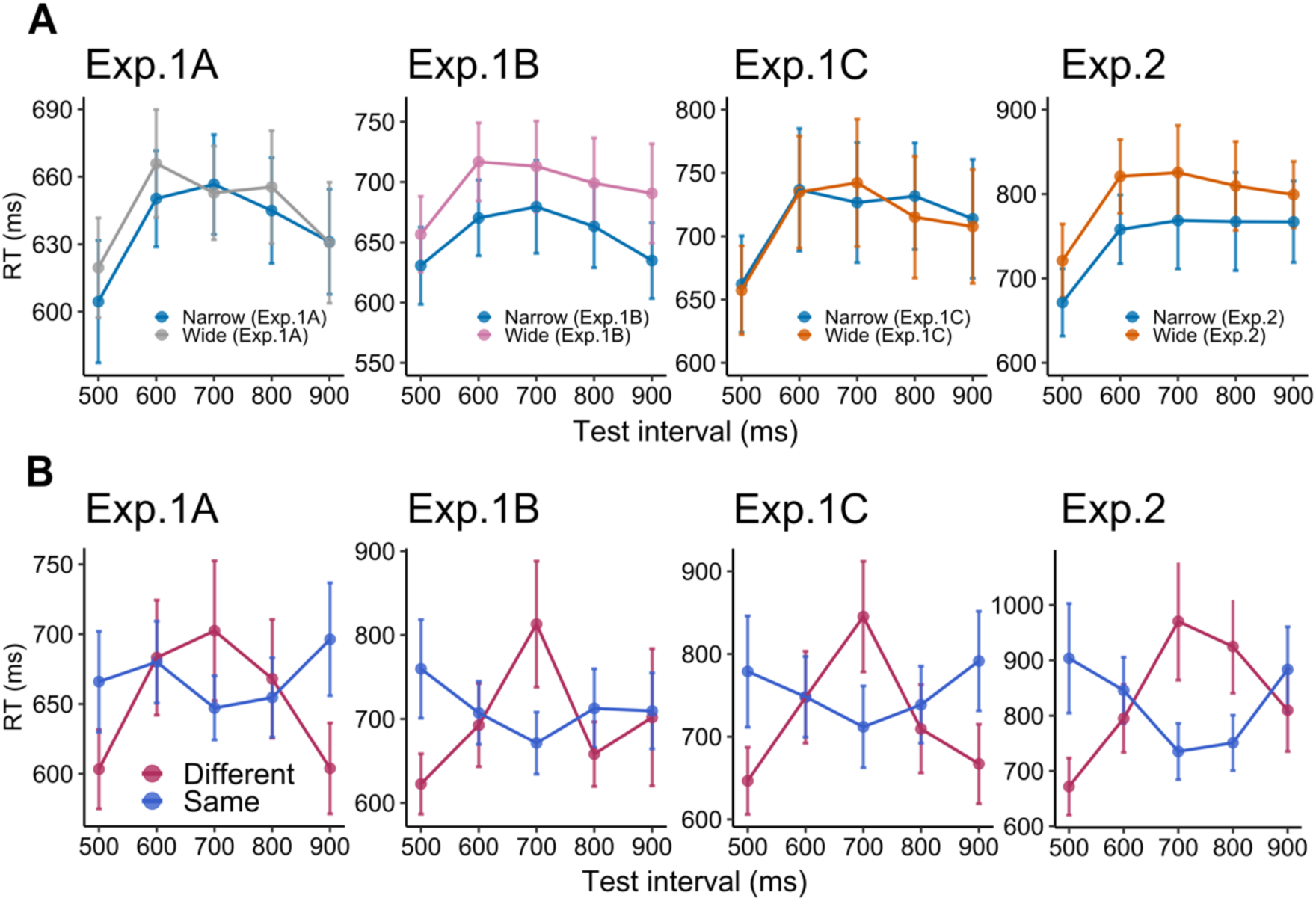
Across-participant mean RTs in the testing phase (temporal generalization task) for each experiment. (A) RT as a function of test interval and distribution. (B) RT as a function of test interval and response (magenta, “different”; blue, “same”; the narrow and wide distributions are concatenated). Error bars indicate SEM.

## Data Availability Statement

The behavioral and EEG data, as well as the experiment and data analysis scripts, are available in the Open Science Framework repository: https://osf.io/3qazh/

## Author Contributions

Taku Otsuka: Conceptualization; Methodology; Investigation; Data Curation; Formal analysis; Visualization; Writing—Original Draft; Writing - Review & Editing. Funding acquisition. Hakan Karsilar: Conceptualization; Methodology; Writing—Review & Editing. Hedderik van Rijn: Conceptualization; Methodology; Resources; Writing— Review & Editing; Supervision; Project administration; Funding acquisition.

## Acknowledgments

We thank Wouter Kruijne and Joost de Jong for their fruitful comments and suggestions on the research design.

